# Brain tumor induce immunoregulatory dendritic cells in tumor draining lymph nodes that can be targeted by OX40

**DOI:** 10.1101/2024.04.01.587531

**Authors:** Oscar Badillo-Godinez, Liam Helfridsson, Jenni Niemi, Shokoufeh Karimi, Mohanraj Ramachandran, Mats Hellström

## Abstract

Brain tumors and metastases have a poor prognosis due to the unique characteristics of the central nervous system (CNS) and tumor immune microenvironment (TIME). CNS tumors exhibit limited infiltration and activation of dendritic cells (DCs) in tumor tissue and tumor-draining lymph nodes (TdLN), which regulate immune responses influenced by factors in the TIME. The immune response in the brain is significantly different from the rest of the body, and although DC subtypes have been identified in mice and humans with brain tumors or metastases, little is known how they affect the response to immunotherapy. We investigated the immunoregulatory function of cervical DCs (DC-c) compared to peripheral DCs (DC-p) in TdLN. Our analysis revealed that DC-c have unique phenotypes and promoted regulatory T cell expansion and poorly cytotoxic CD8 T cells compared to DC-p. Furthermore, we identified OX40 as a modulator of immunoregulatory DC-c function, and *Batf3* knockout confirmed the essential role of DC-c in mounting an immune response to brain tumors. Additionally, the expression of markers associated with mature regulatory DCs (mregDC) in TdLN was associated with immune regulation in the CNS and the response to OX40. Our findings highlight that immunotherapy interventions can modulate DC-c’s immunoregulatory function, offering an innovative approach for optimized immunotherapy against CNS malignancies.

## Introduction

Primary or metastatic cancers in the CNS comprise a diverse group of neoplasms, associated with a poor prognosis irrespective of the cancer type^1^. This unfavorable outlook can be attributed to limited space within the CNS, the detrimental impact of cancer growth, the restricted accessibility of chemotherapy due to the blood-brain barrier, and the unique characteristics of the immune response in the brain^2^. While immunotherapeutic strategies such as Immune Checkpoint Inhibitors (ICIs), oncolytic viruses, and adoptive cell therapies have been extensively employed in extracranial malignancies over the years^3^, their efficacy in CNS tumors has been less pronounced^4,5^. DCs play a crucial role in eliciting specific T-cell-mediated antitumor responses by cross-presenting antigens^6^. In recent years, DC vaccination has emerged as a potential strategy to augment anti-tumor immunity in primary CNS malignancies^7^. In mice, DCs can be categorized into conventional DCs (cDCs) and plasmacytoid DCs (pDCs). cDCs specialize in antigen uptake and presentation to naïve T cells and can be further divided into cDC1 and cDC2, expressing CD8a+ CD103+ and CD11b+ markers, respectively^8^. cDC1 exhibit remarkable abilities in processing and cross-presenting antigens on MHC-I, thereby inducing the polarization of CD8+ T cells, while cDC2 are effective in activating CD4+ T cell responses^7,9–11^. On the other hand, pDCs are the primary producers of type I interferons and play a role mainly in antiviral and antitumor immune responses^12^. Recent Single-cell RNA-sequencing (scRNA-seq) analysis has uncovered additional complexity in DC heterogeneity, identifying multiple subsets such as DC3, which exhibit developmental origin and functional properties associated with an inflammatory gene signature^13^. Furthermore, monocyte-derived DCs (moDCs) represent a subset of DCs that differentiate in response to inflammatory stimuli and are recruited to inflammatory sites^14^. Additionally, within the TIME, cDCs have been found to acquire a common gene program characterized by the expression of immunoregulatory genes (e.g., CD274/PD-L1/PD-L2, and CD200) along with CD40, CD80, LAMP3, and CCR7, leading to the designation of a subset known as mregDCs^15^. DCs are not typically present in the normal brain parenchyma but are primarily found in vascular-rich compartments like the choroid plexus and meninges^16–18^. However, under pathological conditions such as chronic inflammatory diseases, acute infections, neurodegeneration, and cancer, DCs have been observed to migrate to the brain and spinal cord via afferent lymphatics or high endothelial venules^19^. The specific role of DCs in the context of CNS malignancies and their complex interplay with microglia, macrophages, and T cells is still under investigation. One proposed role for DCs is the recognition and presentation of tumor antigens either within the brain or in TdLN to initiate coordinated T cell-mediated responses^20^. However, the TIME has been suggested to promote a tolerogenic phenotype with low expression of costimulatory molecules^21,22^. scRNA-seq analysis of brain and leptomeningeal metastases of melanoma has revealed the presence of various DC subsets, and their proportions have been associated with different responses^23^. Glioma-associated DCs have been linked to a restricted ability to present antigens to T cells, showing an immature cellular state and fostering a tolerogenic phenotype characterized by low expression of costimulatory molecules. Additionally, the use of PD-1 therapy induces changes in costimulatory molecules but fails to overcome the immunosuppressive response^24,25^. Currently, it remains unknown if DCs are necessary or correlate with immunotherapy efficacy in the brain, and the specific DC subtypes present in TdLN of brain tumors and how immunotherapy affects them are yet to be determined.

In this study, we aimed to address these questions by characterizing the phenotype, subtypes, and function of DC-c compared to DC-p in TdLN using two different tumor models representing glioma and CNS lymphoma. Our findings revealed that DC-c exhibited altered distribution of dendric subtypes, similar activation markers compared to DC-p but lower levels of regulatory markers and reduced antigen presentation. Comparing DC subtypes between TdLN from subcutaneous and intracranial tumors showed different gene expression. Functionally, DC-c promoted the induction of Tregs and exhibited lower cytotoxicity and progenitor-exhausted T cells. Furthermore, we demonstrated that the immunoregulatory properties of DC-c could be reversed through αOX40 immunotherapy. Additionally, we identified a correlation between DC markers associated with the mregDC subpopulation and immune regulation within the CNS, contributing to a therapeutic response that dependent on *Baft3-* and LN trafficking.

In summary, our study provides a comprehensive characterization of the immunoregulatory properties of DC subtypes in the TdLN of the brain and elucidates how these properties may vary based on tumor type and therapy.

## Results

### Tumor-draining lymph nodes exhibit distinct immunophenotypes depending on the tumor site

We have previously shown that TdLN from subcutaneous (TdLN-p) tumors exhibit a distinct phenotype compared to those from intracranial (TdLN-c) tumors^26^. Specifically, there are fewer cross-presenting dendritic cells, less active T cells, and more Tregs after immune checkpoint blockade. To further investigate T cell responses, we characterized memory, activation, and regulatory T cell populations from TdLN-c or-p in mice carrying the SB28-OVA glioma model at day 12 post inoculation. (**Figure 1A**). Visualization of the flow cytometry data in a T-distributed stochastic neighbor embedding (t-SNE) map revealed a distinct distribution between naïve lymph nodes (LN) and TdLN, where TdLN-c displayed an accumulation of regulatory and exhaustive markers such as Foxp3, PD-1, and TIM-3 (**Figure 1B**). The analysis of individual populations indicated an increase in the percentage of regulatory T cells in TdLN-c (**Figure 1C**), whereas TdLN-p exhibited a high percentage of Central memory (CD62L+ CD44+) and Effector memory (CD44+ CD62L-)responses, especially in CD8+ T cells (**Figure 1D**). To characterize the memory response in chronic scenarios, we evaluated the expression of killer cell lectin-like receptor G1 (KLRG1), which is expressed in a population that has lost the ability to proliferate but maintains certain functions^27^. TdLN-p showed a high expression of long-term memory in CD8+ T cells (CD44+ CD127+ KLRG1-) compared to TdLN-c, which exhibited a short-term memory profile (CD44+ CD127+ KLRG1+). Moreover, both TdLN showed the expression of markers such as PD-1 and TIM-3 (**Figure 1D**).

**Figure 1.-.**
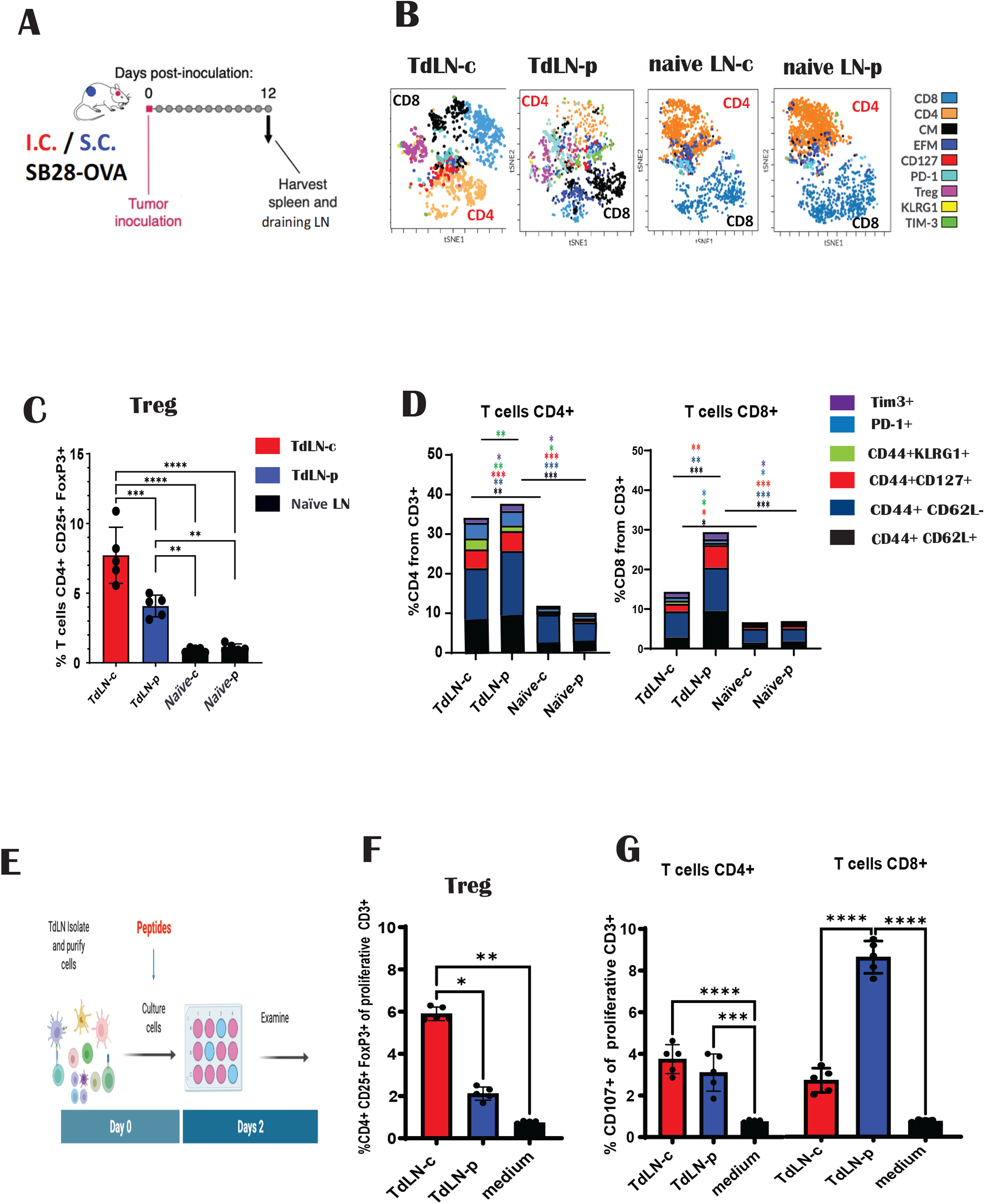
Tumor site-dependent immunological profile of TdLN. Schematic representation of the experiment: Mice were inoculated with SB28-OVA intracranially (i.c.) or subcutaneously (s.c.). On day 12 post-tumor inoculation, TdLN were harvested **(A).** A representative t-SNE map distribution illustrates T-cell populations from TdLN and tumor-free LN (naïve) **(B).** Quantification of regulatory T cells (Tregs) is presented as the percentage of CD4+ CD25+ FOXp3+ cells **(C).** Memory T cell subsets are defined as Effector Memory (CD44+ CD62L-), Central Memory (CD44+ CD62L+), Long-Term Memory (CD44+ CD127+ KLRG1-), Short-Term Memory (CD44+ CD127+ KLRG1+), and Exhausted T cells are identified by the expression of PD-1/TIM3 **(D).** Schematic illustration of the experiment **(E):** Mice were inoculated with SB28-OVA i.c. or s.c., and TdLN were harvested on day 12 post-tumor inoculation. Bulk cells were stained with CFSE and cultured in the presence of OVA peptides for 48 hours. Proliferative cells were analyzed. Tregs (percentage CD4+ CD25+ FOXp3+) **(F)** and cytotoxic T cells (CD107+) **(G)** were evaluated. Graphs display mean ± SEM; analyzed using two-way ANOVA with Tukey’s correction.

To confirm that this profile is not dependent on antigen load, we harvested the TdLN at day 12 post-tumor injection, and the bulk cells were re-stimulated with the same concentration of antigen (**Figure 1E**). We found that under the same conditions, TdLN-c maintained the regulatory response, inducing a higher percentage of Tregs (**Figure 1F**). In contrast, the T cell profile in TdLN-p was less regulatory with a high cytotoxic profile (**Figure 1G**). Simultaneously, we observed an increase in the exhausting profile in TdLN-p, reflecting the activation of the immune response (**Suppl. Figure 1C**).

Taken together, these results indicate that TdLN-c tends to have a T cell response with a regulatory profile. Conversely, TdLN-p possesses a high T cell memory response with a long-term profile, particularly in CD8+ T cells.

### Cervical TdLN DCs exhibit similar profile regardless of brain tumor type

It has been demonstrated that DCs play a crucial role in the response to ICI in various tumors, underscoring their specialized ability to prime CD8+ T cell responses^28^. However, the diversity and functionality of DC-c and how they are influenced by the TIME in CNS are not well understood. We aimed to investigate whether differences exist in the DC population in TdLN. Utilizing a previously reported gating strategy^28,29^, we assessed MHCII+, CD11c+, Ly6c- cells. DCs were distinguished from macrophages based on CD24 high and F4/80 low expression. Subsequently, DCs were categorized into two populations, DC1 and DC2, based on the differential expression of CD8α and CD11b, respectively (**Suppl. Figure 2A**). Additionally, we included migratory DC1, distinguishable as CD103+ or resident DC1 CD103- in the LN^30^. We observed a shift in the percentage from DC1 to DC2 from TdLN-p to TdLN-c, where CD8+ CD103+ in TdLN-p was higher (**Figure 2A**). To determine if this response is specific to the SB28-OVA GBM model or a characteristic of cervical LN, we utilized a lymphoma model (EG7-OVA) to compare immune responses, revealing a similar profile with an increase in DC1 in TdLN-p (**Figure 2B**). At baseline, naïve DCs exhibit low expression of activation markers, maintaining immune tolerance. Both cervical (DC-c) and peripheral (DC-p) TdLN DCs showed essentially the same increase in the expression of activation markers (CD80, CD86, and CD40) compared to naïve DCs. Only CD40 on DC2 from TdLN-p showed a significant increase when compared to DC-c (**Suppl. Figure 2B**). To determine if this profile is specific to TdLN or a systemic response, we examined the profile in the spleen. We found minimal changes in splenic DCs, suggesting that tumor response is more prominent in TdLN (**Suppl. Figure 2C, D**). Furthermore, we assessed antigen presentation by analyzing the expression of the SIINFEKEL peptide-bound MHC-I OVA (H-2Kb OVA257–264). We could detect antigen presentation for both the CD8+ CD103+ and CD103-populations (**Figure 2C, E**). However, both tumor models showed around 50% lower antigen presentation in DC-c compared to DC-p. To evaluate T cell antigen-specific responses, we examined MHC-OVA Tetramers. We observed that in both tumor models, the percentage of Tregs was higher in the TdLN-c (**Figure 2D, F**). Moreover, EG7-OVA showed a memory response profile similar to the SB28-OVA model (**Suppl. Figure 2E**), reinforcing that LN-c promotes a poor cytotoxic T cell response with a regulatory profile.

**Figure 2.-.**
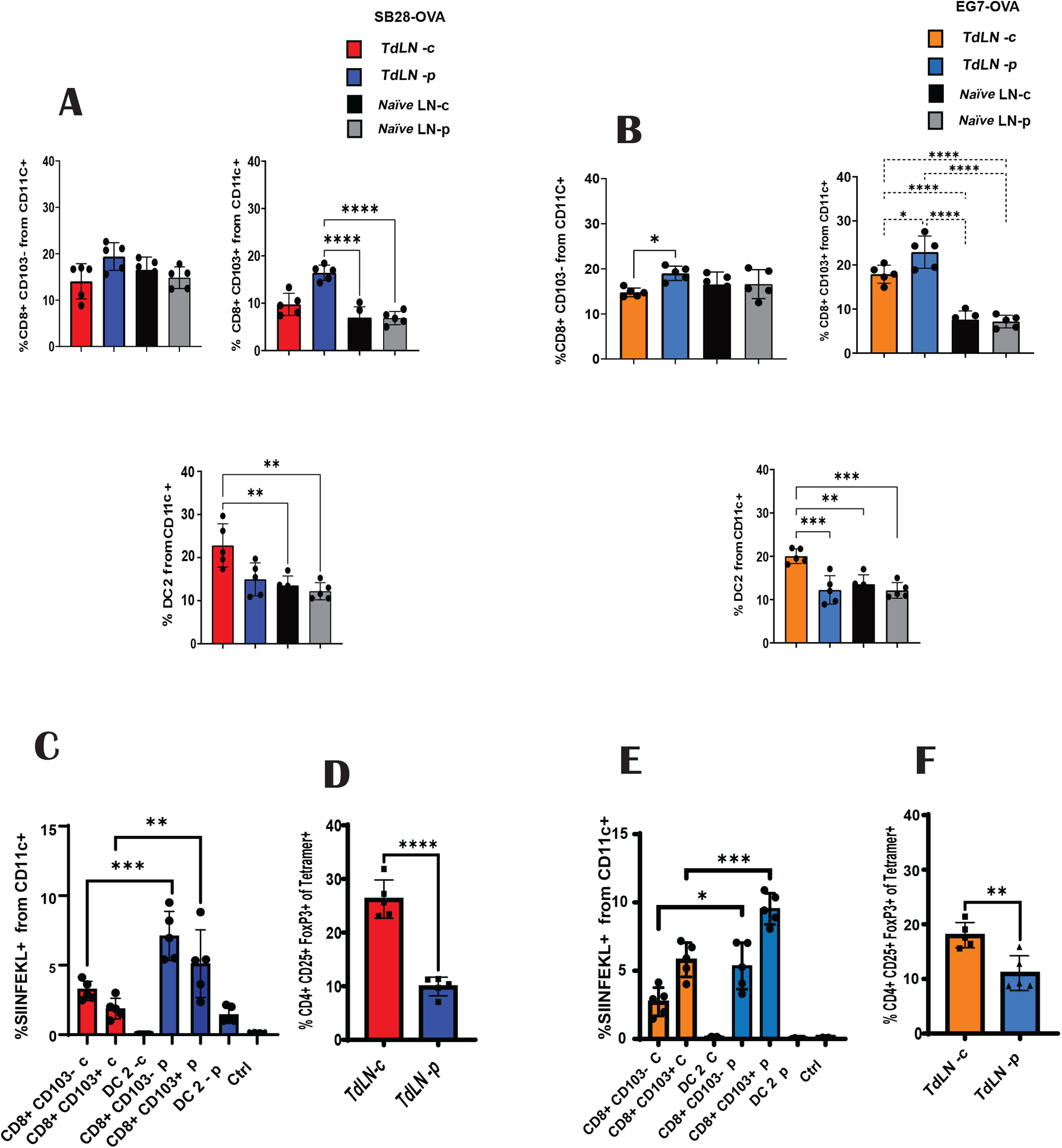
Cervical TdLN have fewer antigen-presenting DCs with tolerogenic function regardless of brain tumor type. The mice were inoculated as in figure 1. TdLN at day 12 post inoculation were harvested. Quantification of DCs subtypes: DC1 (percentage CD8+ CD103+ or CD8+ CD103-), DC2 (CD11b+ CD103-) in SB28-OVA tumors **(A)** or EG7-OVA tumors models **(B).** Panel **(C)** and **(E)** display the percentages of DCs SIINFEKL+ in SB28-OVA and EG7-OVA tumors models, respectively. Panels **(D)** and **(F)** present the quantification of Tetramer OVA+ regulatory T cells (CD4+ CD25+ FOXp3+) in SB28-OVA and EG7-OVA tumors models, respectively. Graphs display mean ± SEM; analyzed using two-way ANOVA with Tukey’s correction, or unpaired t test.

In conclusion, DCs in TdLN-c and TdLN-p do exhibit induction of activation markers, and DC-c exhibits lower antigen presentation and a more prominent induction of a Treg profile compared to TdLN-p.

### DCs from cervical TdLN drives a regulatory T cell response

To functionally study the DC populations, we enriched the DCs (CD11c+ high, MHC II+ high, F4/80-, CD19-) through negative selection and co-cultured them with CFSCS-stained OT I and OT II cells (**Figure 3A**). We observed that TdLN DC-c have a greater propensity to induce Tregs compared to the naïve control and TdLN DC-p (**Figure 3B**). DCs-c showed a significant induction of cytotoxic CD8 T cells, but only half of what DC-p stimulated (**Figure 3C**). When compared to the EG7-OVA model, we observed a similar response trend, with DCs-c inducing a tolerant response and a poor cytotoxic T cell response (**Figure 3 D, E**). However, this regulatory profile was more prominent in SB28-OVA. To confirm that this effect is attributed to DCs, we employed *Batf3* -deficient mice lacking DC1 and treated them in the same manner. The *Batf3* knockout cultures did not show T cell response even when compared with naïve DCs (**Suppl. Figure 3B**). To determine the dominance of each DC population concerning the TdLN site and the type of response, we harvested TdLN DCs from s.c. or i.c. tumors, mixed them in defined ratios (10:1, 1:1 or 1:10), and then co-cultured them with OT-cells (**Suppl. Figure 3C**). The DC-c demonstrated an expansion of Treg and a decrease in cytotoxic T CD8+ profile, with the regulatory profile being more prominent in SB28-OVA compared with EG7-OVA (**Suppl. Figure 3 D, E**). This indicated a dominant effect of the DC-c on the T-cell phenotype, as the DC-p could not significantly counteract the Treg formation induced by DC-c, while DC-c could counteract the development of the CD8 cytotoxic profile. However, we also evaluated the capacity of DCs from TdLN to generate progenitor-exhausted T cells. There was a significant dose-dependent increase in the CD8+ TCF1+ PD1+ T cells driven by DC-p, while DC-c failed to induce TCF1+ expression (**Suppl. Figure 3F, G**).

**Figure 3.-.**
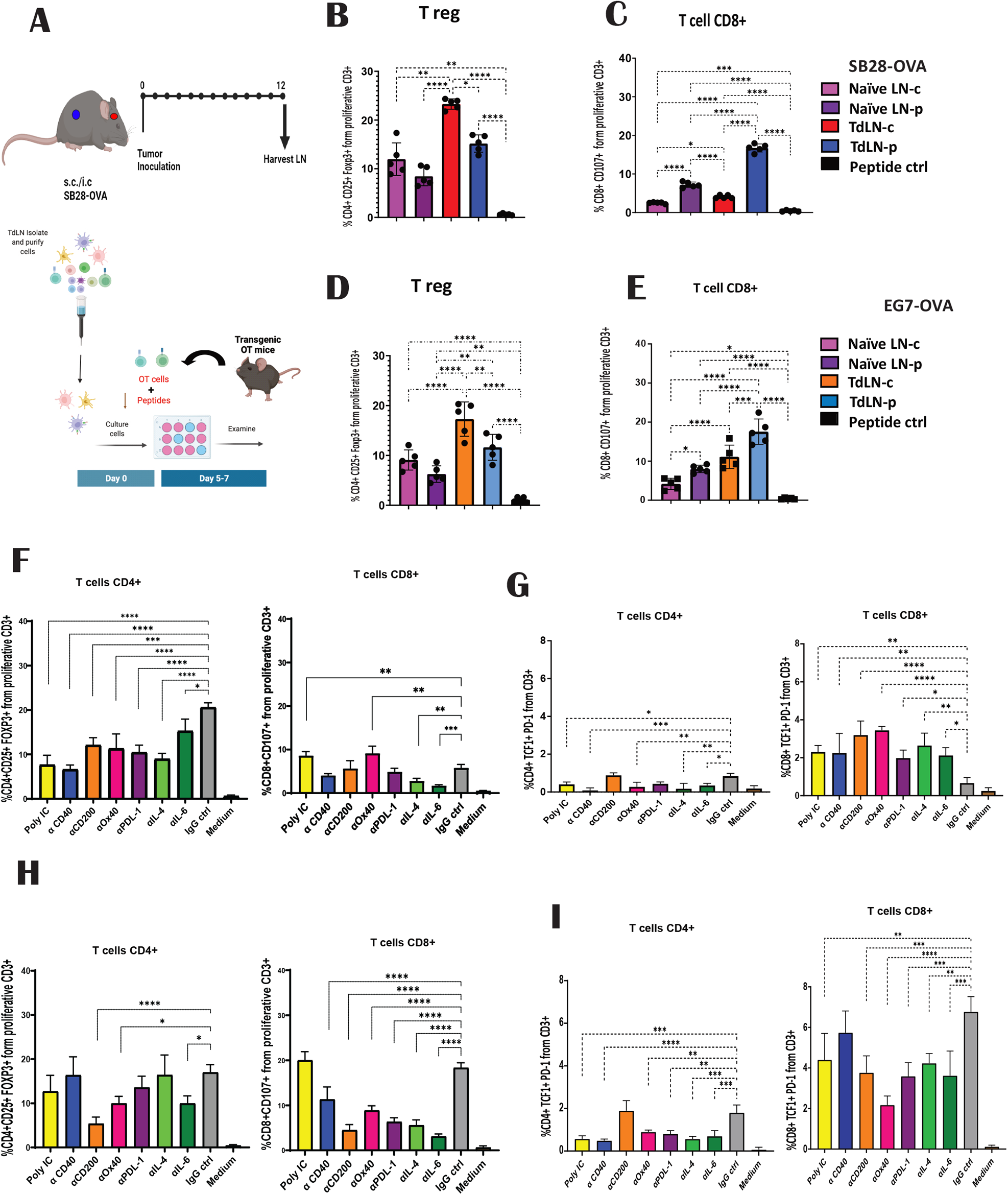
Reprogramming of TdLN DCs Enhances T Cell Response. Schematic representation of experimental procedures **(A):** For *ex-vivo* culture, mice were inoculated i.c. or s.c. On day 12 post-tumor inoculation, TdLN were harvested, and DCs were enriched and co-cultured with pre-stained OT cells in the presence of OVA peptide. The percentage of Tregs (CD4+ CD25+ Foxp3+) and Cytotoxic T cells CD8+ (CD107+) on SB28-OVA **(B, C)** or EG7-OVA **(D, E)** were calculated. Different stimuli were evaluated on *ex-vivo* culture, and the percentage of Tregs, cytotoxic T cell CD8+, progenitor exhausting (Tcf1+ PD-+1) T cell CD4+ and CD8+ from TdLN-c **(F, G)** or TdLN-P **(H, I)** were calculated. Unspecific peptide was used as a control. (n = 5–7 mice/group). Statistics: two-way ANOVA with Tukey’s correction.

Given the tolerogenic profile of DCs from TdLN-c, we sought to enhance their functionality by exploring various pathways through interactions with DC receptors. DCs from TdLN from i.c. and s.c. tumors were isolated and cultivated as described above. While all stimulations showed an effect on downregulating the Treg population, only αOX40 and poly I:C induced robust activation of the cytotoxic profile and progenitor-exhausted CD8+T cells in DC-c (**Figure 3F, G**). For DCs from TdLN-p, only αOX40, together with αCD200 and αIL-6, showed a decrease in Tregs. However, the impact of OX40 on the reduction of cytotoxic and progenitor-exhausted T cells was observed (**Figure 3H, I**).

Taken together, these findings suggest that DCs can exhibit diverse responses depending on the TIME. DC-c play a dominant role in inducing a regulatory T cell profile, thereby hindering the cytotoxic response. Conversely, DC-p demonstrates more efficient activation of T cells, thereby promoting the generation of progenitor-exhausted PD-1+ TCF-1+ T cells. However, this effect can be reversed in DC-c by OX40 stimulation.

### Reprogramming the DC Profile in TdLN Enhances the Immune Response

Consistent with the *ex vivo* experiments, we observed that αOX40 treatment impacted the immune profile of TdLN-c at day 12 post-tumor inoculation (**Figure 4A**). αOX40 significantly enhanced antigen presentation on DC-c CD8+ CD103- and CD8+ CD103+ subsets (**Figure 4B**). Moreover, the therapy induced an increase in CD40 expression on both DC1 subsets, CD8+ CD103- and CD8+ CD103+, while there was no change in the expression of CD86 compared to the control group (**Figure 4C**). Conversely, there were no activation changes observed in DC2. However, the T cell profile was impacted by αOX40, leading to a prominent decrease in the Treg population (**Figure 4D**). An increase in the effector memory (CD62L-CD44+) response (**Figure 4E**) with a long-term (CD127+ KLRG1-) profile was also evident, especially in CD8+ T cells (**Suppl Figure 4B**). Interestingly, the memory T cells exhibited an exhausting profile with characteristics of progenitor exhaustion (**Suppl. Figure 4C, D**).

**Figure 4.-.**
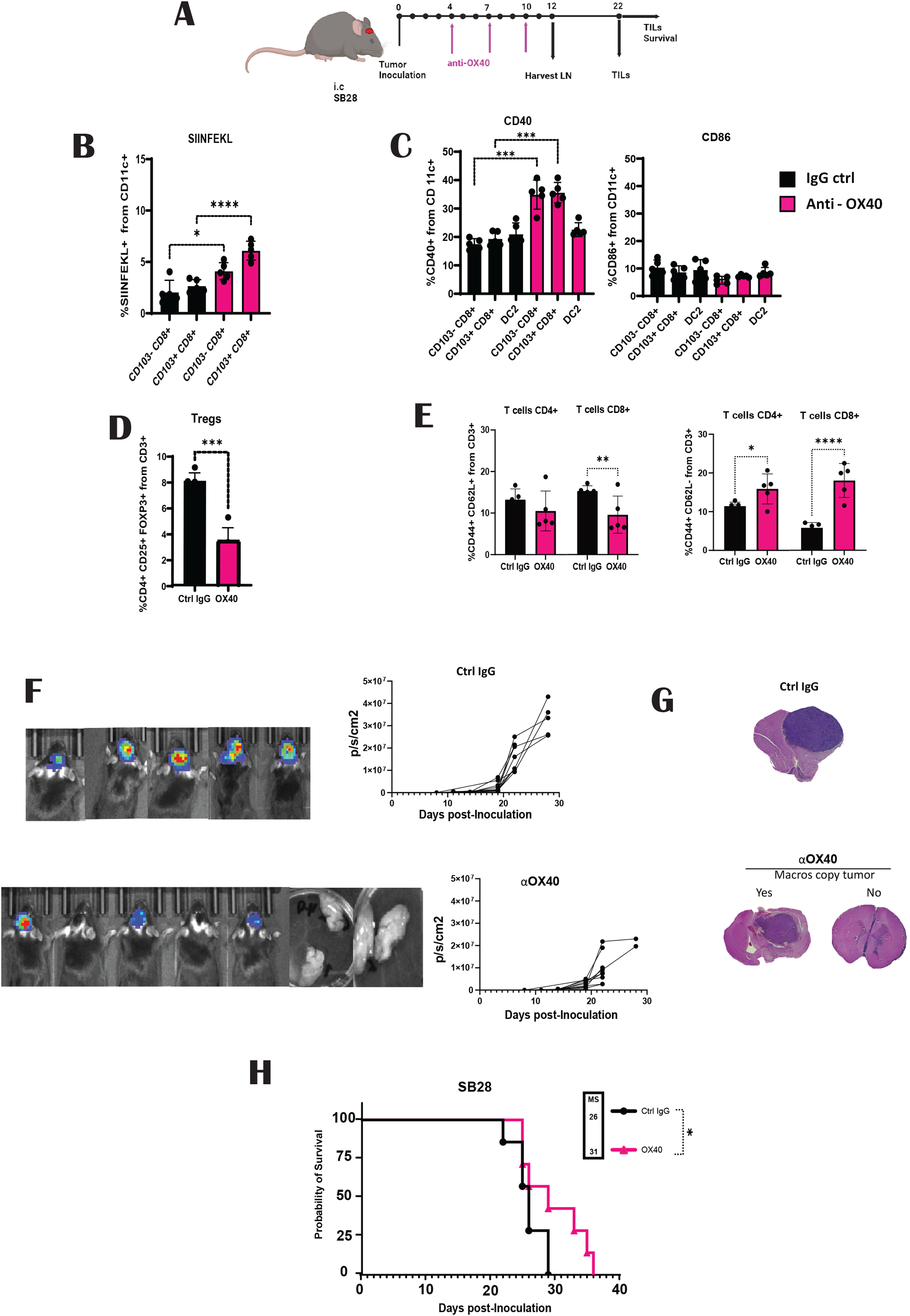
Targeting Anti-OX40 Enhances Immune Responses in Cervical Lymph Nodes. Schematic illustration of the experimental layout (**A**) used to obtain data shown in (**B**)–(**H**): Mice were inoculated with the glioma cell line SB28-OVA, followed by i.p. administration of either αOX40 or IgG Ctrl on days 4, 7, and 10 post-tumor inoculations. On day 12 post-tumor inoculation, the TdLN-c were harvested, and single-cell suspensions were collected for immune profile analysis by flow cytometry. Antigen presentation was analyzed by H2b-SIINFEKL-positive cells on DCs (B). Activation markers CD40 and CD86 from DCs (**C**). Regulatory T cell (CD4+ CD25+ Foxp3+) (**D**). Memory T cell response, for CD4+ and CD8+ central memory (CM, CD44+ CD62L+) and effector memory (EFM, CD44+ CD62L-) from CD3+ cells (**E**). Tumor growth was evaluated by bioluminescence imaging (BLI) (**F**). H&E staining of brain tissue from endpoint (**G**). Kaplan-Meier survival curves of tumor-bearing mice treated with αOX40 or control IgG (**H**). The graph shows the combined survival curves from two independent experiments. Bar graphs show mean ± SEM. One-way ANOVA multiple comparisons test *P < 0.05, **P < 0.01, ***P < 0.001, ****P < 0.0001.

Next, to evaluate how the changes to DC and T cell profiles translated into effects on tumor growth *in vivo*, we performed orthotopic tumor growth studies in mice using SB28-parental. Mice exhibiting a positive IVIS signal at day 10 were selected for the experiment. There was a 50% reduction in bioluminescence reflecting tumor growth in the treated mice (**Figure 4F, G**), resulting in a significant extension in median survival of 31 days compared to the control group’s 26 days (**Figure 4H**).In parallel, we analyzed tumor-infiltrating lymphocytes (TILs) on day 22 post-inoculation, just before the mice displayed symptoms. The fraction of T cells associated with cytotoxic potential (CD107+) was lower in the control group than in the αOX40-treated group.

This difference was more pronounced in CD8+ T cells, regardless of whether the mice exhibited macroscopic tumors or not. Additionally, T cells demonstrated a proliferative profile, especially in CD4+ T cells, in mice with no macroscopic tumors compared to the control group. Furthermore, mice treated with αOX40 showed a significant difference in the progenitor exhaustion profile. Notably, regulatory T cells were significantly fewer in αOX40-treated mice than in their counterparts (**Suppl. Figure 4E**). Analysis of TILs at the endpoint from mice treated with control IgG and αOX40 showed that they maintained a cytotoxic response, along with a proliferative and exhausting progenitor profile, regardless of tumor growth. Additionally, T cells maintained a lower Treg profile at the endpoint stage (**Suppl Figure 4F).**

The survival benefit was small given the strong effect on tumor bioluminescence signal and low tumor burden upon post-mortem examination in the SB28-parental mice. We hypothesized that an activation of the immune response following therapy could reduce survival in a fraction of the OX40-treated mice by induction of necrosis and edema. Therefore, we explored the use of low-dose dexamethasone, a potent corticosteroid administered to brain tumor patients to reduce tumor-associated edema^31^. The combination treatment with low-dose dexamethasone administered to all mice on day 22 significantly increased median survival (42 days), resulting in a survival rate of 12.5%, with no effect in the control-treated group (**Suppl. Figure 4G, H**).

Collectively, these findings demonstrate that αOX40 has the potential to reprogram the TIME and DC-c, enhancing antigen presentation and activation profiles, thereby improving the T cell profile in TdLN and leading to a therapeutic response in the brain.

### DCs subpopulations from cervical TdLN nodes exhibit distinct RNA expression profiles

Given the distinct functional differences between DC-c and DC-p, we conducted a comprehensive analysis of DCs subpopulations using combined antibody and targeted scRNA-seq to profile MHCII+ CD11c+ cells from TdLN-c and TdLN-p. Our analysis, performed on both naïve and tumor-bearing TdLN samples, revealed six distinct clusters (**Figure 5A**). Among these clusters, the DC1 cluster included genes *Cd8a* and the CD103 marker, while the DC2 cluster comprised *Itgam* alongside the CD11b marker. The pDCs cluster showed *Klra17, Fcer1g, Irf8*, and *Tlr7*, while the monocyte-derived DCs (moDC) cluster included *Cd9, Arg1*, and *Itgam* (**Suppl. Table 3**). Additionally, we identified the DC3 cluster expressing *Xcr1, Ifngr1, Cd86*, and *Tlr3*, which was distinct from the mregDC cluster. The mregDC cluster exhibited expression of maturation markers such as *Cd40, Relb*, and *Lamp3*, along with immunoregulatory genes like *Cd274, Pdcd1lg2, Fas, Socs1,* and *Aldh1a2* (**Figure 5B**).

**Figure 5.-.**
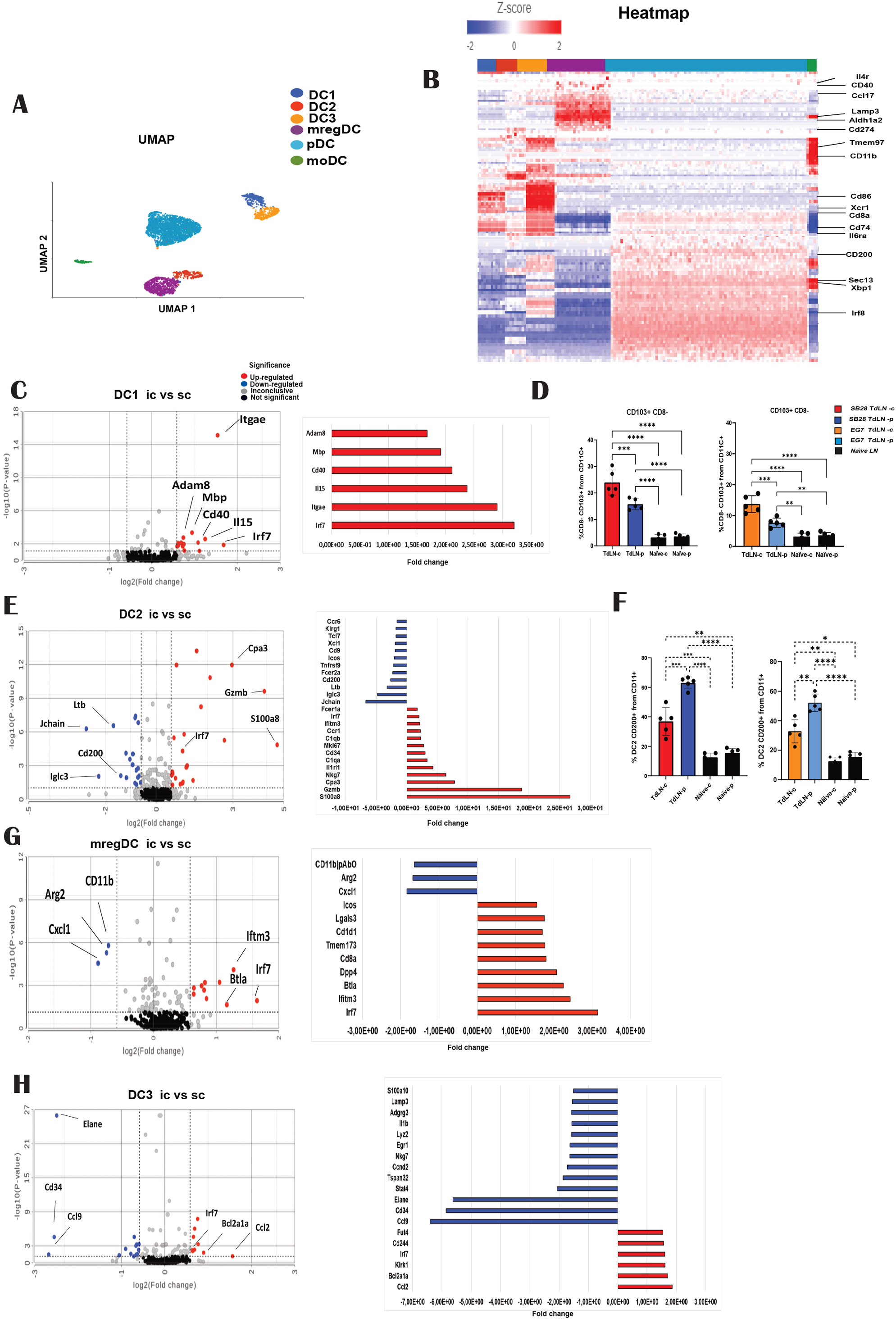
mregDC Profile in Cervical Lymph Nodes Correlates with *α*OX40 Immunotherapy. SB28-OVA or EG7-OVA were implanted i.c. Mice were sacrificed on day 12 post-tumor inoculation, and the TdLN were collected for scRNA-seq analysis or flow cytometry analysis. Uniform manifold approximation and projection (UMAP) of scRNA-seq profiles of 9,904 TdLN cells (**A**) was conducted. Heatmap of genes upregulated in each DC cluster was generated, with values presented as averaged Z scores (**B**). Statistical analysis involved filtering genes based on a false discovery rate (FDR) cutoff of <0.05. FDR (Benjamini–Hochberg adjustment) was calculated from p-values obtained through ANOVA. Differential gene expression associated with TdLN-c versus TdLN-p on DC1 (**C**) DC2 (**E**), mregDC (**G**) and DC3 (**H**). Calculation of DCs CD103+ CD8- (**D**) and DC2 CD200+ **(E)** from TdLN SB28-OVA or EG7-OVA by cytometry. Bar graphs show mean ± SEM, two-way ANOVA with Tukey’s correction: *p < 0.05, **p < 0.01, ***p < 0.001, ****p < 0.0001.

Next, we conducted Gene Set Enrichment Analysis (GSEA) and Ingenuity Pathway Analysis on DC1, DC2, DC3, and mregDC clusters, revealing that DCs-p exhibited significant representation of multiple genes involved in immune response pathways. These pathways included: adaptive immune response, regulatory immune response, positive immune effector function, cytokine production, IFN response, and proliferation and activation of Th1 and Th2 (**Supp Figure 5A, B, Suppl. Table 4**). Upon intra-cluster analysis, DC1-c displayed upregulation primarily in *Irf7* and *Itgae* (**Figure 5C**). To further investigate the expression of *Itgae* on the subpopulation of DC1, we employed a flow cytometry–based strategy, focusing on the CD103+ CD8-population, which demonstrated a higher percentage in TdLN-c compared to TdLN-p (**Figure 5D**). Moreover, DC2-c cluster exhibited high expression of *S100a8, Gzmb*, and *Cpa3,* along with downregulation of *Jchain, Iglc3, Ltb*, and *Cd200* (**Figure 5E**). Consistent with RNA expression, cytometry analysis revealed low CD200 expression in DC2-c compared to DC-p (**Figure 5F**). Further, the mregDC-c had upregulation primarily in *Irf7, Ifthm3*, and *Btla*, with downregulation observed in *Arg2, Cxcl1*, and *CD11b* (**Figure 5G**). Additionally, DC3-c displayed upregulation of Ccl2 and Bcl2a1a, while showing downregulation mainly in *Elane, Cd34*, and *Ccl9* (**Figure 5H**). Notably, all DCs-c clusters exhibited increase on expression of *Irf7*.

Taken together, these findings highlight gene expression differences between TdLN-c and periphery TdLN DCs subpopulation within the defined subtypes of DCs.

### mregDC Profile is associated with the Impact of OX40 Treatment and depends on Baft3 DCs

Recently mregDCs with characteristic markers such as CD200, CD40 and CCR7 has been identified as a key element in the TIME, contributing to the control of the anti-tumor immune response^15^. We observed that *Cd200* was expressed in all DC populations, while *Cd40* was more pronounced on mregDCs, along with a predominant expression of *Ccr7*. Additionally, we noted a predominant expression of the OX40 gene (Tnfrsf4) on mregDCs (**Figure 6A**). Based on the scRNA seq profiling above, we further assessed the proportion of mregDCs in the TdLN-c using flow cytometry (**Suppl. Figure 5C, D**). The fraction of mregDCs (CD11c+, MHCII+, Ly6c- CD24+, F4/80-, CD40+, CD200+, and CCR7+) was more than double in DC-p irrespective of model and in EG7 DC-c as compared to DC-c from SB28 bearing mice (**Figure 6B**). Consequently, we assessed the anti-tumor response in the EG7 model, characterized by a high proportion of mregDC in the TdLN, using αOX40. To determine if the response relied on DCs, we employed DC-deficient mice with *Batf3* knockout. The absence of *Batf3* DCs resulted in reduced survival in the i.c. EG7 model without any treatment, indicating an essential role of DCs in the normal immune response in this lymphoma model (**Figure 6C**). Subsequently, mice were treated with an OX40 agonist, which elicited a robust response, leading to a 71% survival rate. The treatment effect of αOX40 was completely dependent on *Batf3*, and there was an 8-fold reduction in mregDC upon αOX40 treatment.

**Figure 6.-.**
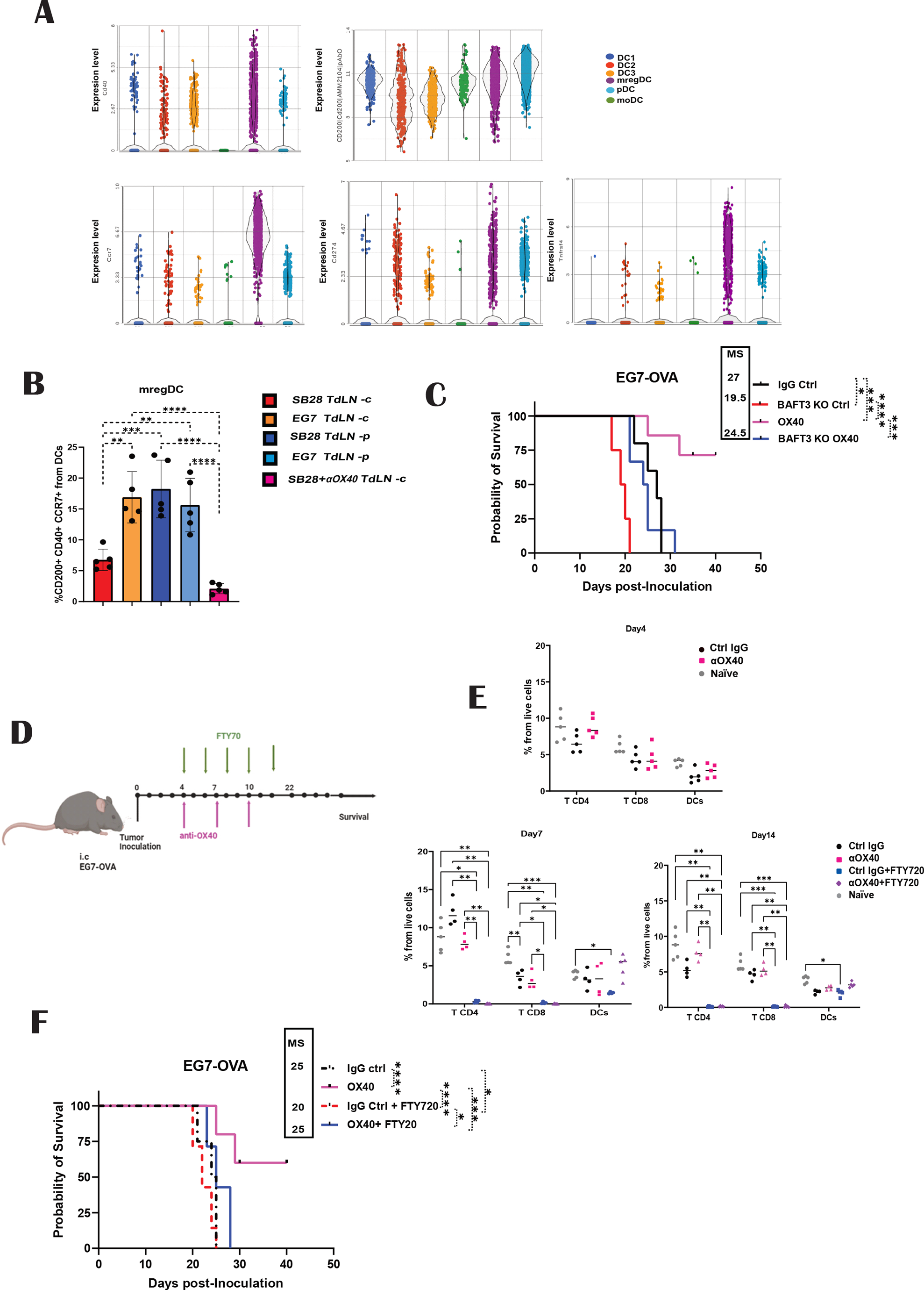
mregDC Profile in Cervical Lymph Nodes Correlates with *α*OX40 Immunotherapy. Gene expression levels of Cd40, CD200, Ccr7 and Tnfrs4 from DCs clusters (**A**). Statistical analysis involved filtering genes based on a false discovery rate (FDR) cutoff of <0.05. FDR was calculated from p-values obtained through ANOVA. The calculation of mregDC (CD11c+ MHCII+ Ly6c- CD24+ F4/80- Ccr7+ CD200+ CD40+) in TdLN was performed by cytometry (**B**). Kaplan-Meier survival curves of EG7-OVA tumor-bearing wild type or Baft3KO mice treated with αOX40 or control IgG (n = 5-7 mice/group) were analyzed using the log-rank test (**C).** Schematic experimental representation **(D).** Kaplan-Meier survival curves of EG7-OVA and tumor-bearing mice treated with either OX40 or IgG control on day 4, 7 and 10 post tumor inoculation. At the same time mice were inoculated i.p with FTY70 starting on day 4 post tumor inoculation and every other day (n = 5-7 mice/group) were analyzed using the log-rank test. Statistical analysis was conducted using the Mann-Whitney test. Bar graphs show mean ± SEM, two-way ANOVA with Tukey’s correction: *p < 0.05, **p < 0.01, ***p < 0.001, ****p < 0.0001.

Finally, to assess the role of TdLN in therapy response, we investigated the impact of systemic administration of FTY720 (**Figure 6D**), which blocks lymphocyte LN trafficking^32^. Circulating T cells and DCs were analyzed 24 hours after drug administration using flow cytometry (**Supp Figure 5E**). FTY720 administration resulted in significant reductions in the frequency of circulating CD3+ T cells in the blood. Interestingly, the effect of FTY720 on DCs was more pronounced in the IgG control group, while OX40 therapy demonstrated a recovery in DC percentage (**Figure 6E**). However, mice injected with FTY720 failed to respond to αOX40 therapy (**Figure 6F**), indicating that lymphocyte trafficking from lymphoid tissues to tumors plays a critical role in tumor eradication.

These findings underscore the crucial role of DCs in orchestrating an immune response against brain tumors. Moreover, they highlight the association between the presence of mregDCs in TdLN-c and the efficacy of αOX40 therapy in enhancing the immune response.

## Discussion

In this study, we demonstrated that TdLN DC-c, which have a unique connection to the brain, are predisposed to induce a tolerogenic function. We employed the SB28 glioblastoma model with a distinct advantage, in comparison to the widely used preclinical GL261 GBM tumor model as it does not respond to ICI, and thereby mimics human GBM^33^. Previously, we showed that the TIME dictates the ICI response to SB28, as i.c. tumors were unresponsive to ICI, while SB28 s.c. tumors showed a positive response, resulting in the development of a memory response against brain tumors^26^. Consequently, we further characterized TdLN-c with TdLN-p to better understand the difference in the immune response. Additionally, we compared these differences using a distinct tumor cell line, EG7-OVA, which is a T-cell lymphoma model. Both models showed that TdLN-c exhibits a tolerogenic immune response, characterized by increased Treg levels and poor activation with low cytotoxic T cell response. Through the enrichment of the DC population and *ex vivo* culture, we demonstrated a direct influence on the activation and generation of regulatory T cell response. Although the enrichment was not 100% pure and remnants of other cells could affect T cell activation, the direct effect from DCs was supported by the use of *Batf3* knockout mice, which are primarily deficient in DC1 with minimal changes in the DC2 population. Consequently, in the absence of TdLN DC1, we observed a decrease in T cell activation, comparable to levels seen with naïve DCs. The relation of DC1 to T cell activation has previously been demonstrated, as it is the primary source of IL-12 and plays a crucial role in both the induction and maintenance of T cells^34^. Early priming of CD4+ T cells against tumor-derived antigens also necessitated DC1, as confirmed by the selective deletion of CD40 and MHC class II molecules in DC1. This selective deletion is critical not only for CD8+ T cell priming but also for the initial activation of CD4+ T cells^35^. Furthermore, *ex vivo* cultures showed the advantage of manipulating the DC profile. We found that different stimuli exhibited varying effects on the activation of T cells. Poly I:C and OX40 induced more cytotoxic profile in TdLN-c, however, our previous findings indicated that Poly I:C does not significantly impact the immune response against SB28^26^. This suggests that stimulating T cells by targeting OX40 could promote strong activation in DCs, thereby enhancing the immune response. This is relevant because OX40, a receptor expressed on T cells, and its ligand OX40L, primarily expressed on DCs, induce T cell proliferation and reduce Tregs development^36^. In the context of CNS malignancies, targeting OX40 has been shown to enhance survival in mice, especially when combined with vaccination using irradiated tumor cells or GM-CSF-expressing GL261 cells (GVAX). This combined approach significantly improves survival compared to vaccination alone. A similar study conducted by Murphy K et al., employing a combination of tumor lysate vaccine, revealed that targeting OX40L has a more pronounced effect against the GL261 glioblastoma mice model^37^. This supports the hypothesis that stimulating the production of DCs population through the addition of GM-CSF and stimulation via OX40L could play a crucial role in the response to CNS malignancies, with the caveat that GL261 is very responsive to ICI. Additionally, it has been shown that targeting OX40 has the potential to decrease IL-10 levels, facilitating the maturation of DCs^36,38,39^. However, neither of these studies explored the relationship between OX40 activation and its importance in the DC immune response. We found that αOX40 treatment increased the percentage of antigen cross-presentation with the overexpression of CD40 in the DC1 subpopulation. Despite the improvement in DC and T cell activation and its effect on tumor growth and median survival, none of the mice survived. The analysis of T cell response in the TIME showed that TILs at different time points from mice with or without macroscopic tumors revealed that OX40 therapy had an impact on Tregs. On the other hand, the progenitor-exhaustive profile was generally increased as well, suggesting that early activation and reinvigoration of T cell activity effectively slowed down tumor growth. The data suggest that the treatments potentially elicit an exaggerated response, leading to side effects, as evidenced by mouse deaths despite successful tumor elimination. Additionally, treated mice had reduced viable tumor volume as compared to control mice. SB28-parental exhibit variable amounts of tumor necrosis and edema (Figure 4G) and therefore we explored if concurrent dexamethasone after the induction of a therapeutic response could mitigated any exaggerated consequences of therapy^40^. Low doses of dexamethasone had a positive effect in combination with αOX40 therapy, resulting in a 12% increase in survival and prolonged median survival time.

Recently, additional DC subpopulations have been identified both intratumorally and in TdLN as crucial elements playing a role in regulating the immune response (DC1, DC2, DC3, mregDC, moDC)^15,41,42^. Utilizing scRNA-seq in human and mouse, several groups have described a conserved program consistently identified in DC subpopulations^6,43^. By employing combined antibody and targeted scRNA sequencing, we aimed to identify the DC population in TdLN from both SB28 and EG7 models. Through the enrichment of DCs and employing Abseq targeting CD11b, CD103, F4/80, and CD200, we identified six major clusters, including the observed mregDC cluster, characterized by an activated and regulatory profile, distinct from the DC3 cluster.

DC1s are well known for their efficient cross-presentation capacity, not only in response to infection but also in anti-tumor immunity^30,44^. Migratory DC CD103+ and lymphoid-resident DCs CD8+ share many attributes, such as dependence on the same transcription factors, cross-presenting ability, and expression of certain surface molecules^45,46^. Despite this, the functional diversity of the two cDC1 types is not fully understood. Reports on LN-resident DCs (CD8+ cDC1) strongly advocate their role as a platform for CD4+ T cells in enhancing memory CD8+ T cell formation, while migratory CD103+ DCs from draining lymph nodes were found to be more potent at inducing Th17 cytokine production by CD4+ T cells than DCs^47^. We did not find a difference in the proportion of CD103+ CD8a+, but CD103+ CD8- showed a higher percentage in cervical LN. Further studies on this subpopulation would be interesting to reveal more information on DC subpopulations and their relation with brain tumors.

A tumor-protective role of cDC2s has been demonstrated in experimental mouse models. CD11b+ DCs that had infiltrated B16 melanoma can prime T cells but were characterized by reduced capacities for antigen uptake, antigen presentation, and migration to TdLNs, compared with normal skin DCs^48^. Additionally, cDC2s were found to promote the growth of MC38 tumor cells and to inhibit the infiltration of Th1 and TNF-α–producing cells into the tumor^49^. These observations suggest that cDC2s, at least in part, attenuate antitumor immunity, possibly by limiting antitumor CD4+ effector T cell responses. In our study, we observed an increase in the DC2 population in TdLN-c. cDC2-c showed an increase in the expression of *S100a8*, a calcium- and zinc-binding protein that plays a prominent role in the regulation of inflammatory processes and immune response^50^. We also observed downregulation of CD200 on cDC2-c. The expression of CD200 in DCs is overexpressed during inflammatory signals such as apoptosis and production of IFN-γ and TNF-α, mediated by CD200R1 engagement^51^. On the other hand, it has been hypothesized that triggering of alternate pathways, particularly CD200R, on DC precursors leads to biased differentiation towards so-called “tolerogenic” DCs^52^. This suggests that DC2 in cervical LN may not response to inflammatory response and have a predominant regulatory response.

We observed upregulation of IRF 7 in all clusters. IRF7, a member of the IRF family of transcription factors, acts as the ‘master regulator’ of type I interferon (IFN)-dependent immune responses, underpinning their critical role in host defense^53^. Furthermore, large scale analysis of genes regulated by IRF7 in response to viral infection have identified CD80 as a potential target of this transcription factor in an *in vitro* system^54^. Reports have shown that IRF7-deficient cDCs exhibit heightened responsiveness to TLR2, characterized by increased CD86 expression both *in vitro* and *in vivo*, as well as altered cytokine profiles that affect their capacity for CD4+ T cell polarization^55^. This implicates IRF7 as a component of a regulatory pathway initiated in dendritic cells during their response.

The difference between the mregDC and DC3 populations has been debatable. The downregulation of canonical cDC1 and cDC2 gene transcripts observed in the identified DC3 cluster aligns with the decreased expression of these genes during dendritic cell maturation^46^. This maturation process involves specialized antigen presentation programs and is associated with an inflammatory gene profile, as documented in previous studies^56,57^. Our findings suggest that mregDCs, also known as CCR7+LAMP3+ cells^42^, represent a distinct molecular state preferentially with cDC1s profile in cervical LN. Moreover, the distinction between mregDCs and DC3s in T cell activation remains unclear, and the specific details are not yet understood. Further experiments should be conducted, and the sample size increased to distinguish subtle changes between DCs subpopulations to understand the molecular system that dynamically regulates their development and maintenance. Here, we focus on the mregDC subpopulation, as previous research has shown that preclinical studies and early-stage clinical trials play critical roles in response to ICI^15,41^. We observed a notable difference in the percentage of mregDCs in TdLN-c, with EG7 showing a higher percentage compared to SB28. Furthermore, EG7 αOX40-treated mice exhibited a robust response against brain tumors with high efficacy. The necessity of *Batf3* DCs for cross-priming CD8+ T cells and generating de novo anti-tumor T cell responses has been demonstrated^58^. We found that *Batf3* knockout mice failed to respond to αOX40 therapy, indicating the relationship between DC1 and mregDC roles in mediating the anti-tumor response. The percentage of mregDCs was strongly downregulated when SB28 was treated with αOX40. This suggests that the impact of OX40 in downregulating Tregs could trigger the activation of DCs, inducing a proinflammatory state and subsequently downregulating the mregDC population. This phenomenon may potentially result in an overreaction in the CNS. Furthermore, the response to immunotherapy αOX40 was dependent on LN trafficking, as blocking with FTY720 negatively impacted survival. This highlights the role of TdLN in mounting the tumor immune response and its interaction with immunotherapy. The importance of mregDCs on T cell activation in TdLN is not yet understood, and the specific details remain unclear, highlighting the need for further research to better comprehend the molecular system dynamically regulating their development and maintenance in the TIME.

In conclusion, DCs from brain TdLN exhibit distinct genetic, phenotypic, and functional characteristics, demonstrating a notable default regulatory program compared to peripheral TdLN DCs. The pivotal role of DCs in antigenic stimulation and activation stands out as a crucial factor for the success of immunotherapy in CNS malignancies. Our findings demonstrate that reprogramming the DC-c profile through modulation of OX40, holds promise for achieving an anti-tumor response and improving survival outcomes. A comprehensive understanding and accurate characterization of the states of tumor DCs will unveil molecular targets with the potential for immunotherapies, refining existing treatments, and designing innovative targeted strategies.

## Methods

### Cell lines

The SB28-parental cells were cultured in DMEM supplemented with 10% Fetal Bovine Serum, 2 mM Glutamine, 100 U/ml penicillin, and 100 mg/ml streptomycin (Gibco™). To establish an SB28 cell line expressing the full-length OVA peptide (SB28-OVA-FL), the pAC-Neo-OVA plasmid from Addgene (#22533) was utilized, and the cells were cultured under selective antibiotic pressure with 400 μg/ml G418 (Gibco™). The EG7-OVA cells were cultured in RPMI with 10% FBS, 50 μM 2-ME, 400 μg/ml G418 (Gibco™), and 2 mM glutamine. All cells underwent authentication and testing, and their mycoplasma status was monitored monthly using the PlasmoTest Mycoplasma Detection Kit (InvivoGen).

### Mouse studies

The animal experiments involved female C57BL/6 mice, aged 6-8 weeks, obtained from Taconic® and acclimated for 7 days prior to injection. Transgenic mice OT-I and OT-II C57BL/6-Tg were acquired from Charles River©. All experiments were conducted following national guidelines and regulations (permit 5.8.18-14723/2020).

#### Orthotopic murine glioma models

SB28 parental or SB28-OVA cell lines were cultured and re-suspended to form either 1600 cells or 2.5 × 10^4^ cells for EG7-OVA to be inoculated intracranially (i.c). The mice were anesthetized with isoflurane and positioned on a stereotactic frame. A burr hole was drilled in the skull using a 0.7 mm drill bit, positioned 1.5 mm laterally anteroposterior and 1.5 mm mediolaterally from the bregma. A non-coring needle (Hamilton 7804-04, 26-gauge, Small Hub RN Needle, Point Style: 4, Needle Length: 2 inches, Angle: 12) was then used to inject the cells at a depth of 3 mm into the brain through the burr hole. The burr hole was covered with bone wax, and the skin incision was sutured. Mice were administered an intraperitoneal injection of carprofen (0.05 mg/kg) immediately after surgery and again 4-6 hours post-surgery, and they were monitored for signs of pain. Only mice showing no signs of pain 24 hours post-surgery were included in the study. Tumor-bearing mice were monitored for tumor growth 1-2 times weekly via bioluminescence imaging using an IVIS Spectrum in vivo imaging system. Prior to imaging, the mice were injected intraperitoneally with 15 mg/kg luciferase (AAT Bioquest® 115144-35-9) and anesthetized with isoflurane. The signal was analyzed using the IVIS® Lumina III PerkinElmer.

### Mouse subcutaneous flank injections

Mice were subcutaneously injected at the right-sided flank region with SB28-OVA or EG7 cells suspended in PBS at a concentration of 5×10^5^ cells. Tumor volume in subcutaneous flank tumor-bearing mice was measured 1-2 times weekly using caliper measurements, and volume was approximated using the modified ellipsoid formula (length x width^2)/2.

For in vivo treatment, mice were intraperitoneally treated with 200 μg of agonist OX40 BioXCell (OX-86) or a control group was treated with 200 μg of BioXCell control IgG on days 4, 7, and 10 post-tumor inoculations. Mice were monitored daily and evaluated for tumor-related symptoms such as hunched posture, lethargy, persistent recumbency, and weight loss using the Uppsala University (Uppsala, Sweden) scoring system for animal welfare. The mice were sacrificed at the humane endpoint, determined when the welfare score reached a maximum of 0.5. Alternatively, mice were sacrificed at the experimental endpoint on day 12 post tumor injection to collect TdLN for ex vivo experiments and single-cell (sc)-RNA sequencing or flow cytometry analysis.

For FTY720 treatment, mice were intraperitoneally injected with 25 µg of FTY720 (APExBIO©) on the same day the treatment began, and then received maintenance doses of 20 µg of FTY720 every other day until day 16 post tumor inoculation, when mice began to show symptoms.

### Harvesting of Lymph Nodes

The mice were euthanized, and the cervical lymph nodes (LN-c) or inguinal lymph nodes (LN-p) were harvested and placed in fresh RPMI media. They were then incubated for 10 minutes at 37°C with Liberase (250 μg/ml) and DNAse (50μg/ml Merck®). After incubation, the lymph nodes were smashed through a 70 μm cell strainer in PBS. Splenocytes were harvested and incubated with RBC lysis buffer (Invitrogen™) for 5 minutes, followed by washing and resuspension in PBS.

#### T-cell Isolation

Splenocytes from OT-mice were resuspended in MACS buffer (PBS, 0.5% BSA, 2mM EDTA) with anti-CD4 Microbeads or anti-CD8 Microbeads (Milteny Biotec ©) according to the supplier’s instructions. They were then incubated for 10 minutes on ice, washed with MACS buffer, and transferred to the magnetic columns for positive collection.

### Dendritic Cell Isolation

Cells from the lymph nodes were resuspended in MACS buffer along with Pan Dendritic Cell Biotin-Antibody Cocktail (MiltenyBiotec ©) following the supplier’s instructions. The cells were then recovered by negative selection.

### Ex Vivo Experiments

The OT cells were incubated with 5μM of green CellTrace™ solution (Thermo Scientific) for 10 minutes at 37°C, then washed and resuspended in tissue culture medium RPMI with 10% FBS, 50 μM 2-ME, 50µ/ml Geneticin, and 2 mM glutamine. T cells were cultured in 96-well plates with 50×10^3^ T cells (1:1 OT-I and OT-II cells) and co-cultured with 5×10^3^ TdLN DCs obtained from either i.c. or s.c. sources. Various stimuli were added to the culture: Poly I:C 5µg/ml (Adipogen Life Sciences), αCD40 (FGK45), αIL-6 (MP5-20F3), αIL-4 (BVD6-24G2), αOX40 (OX-86), αCD200 (OX-90), and αPDL-1 (10F.9G2TM) from BioXcell. In parallel experiments, OT cells were cultured as described above with DCs from TdLN-c mixed in ratios of 1:1, 10:1, or 1:10 with DCs from TdLN-p at a final concentration of 5×10^3^ dendritic cells. OVA peptide H-2kb SIINFEKL, 323–339 OVA I-Ab (Merck©), and control peptides H-2Kb glycoprotein BSSIEFARL (TS-M523-P MBL©) and HBc 128-140 aa TPPAYRPPNAPIL (TS-M701-P MBL©) were added at a concentration of 10 μg/ml and incubated for 5-7 days at 37°C.

### Extraction of TILs

Single cells were obtained from tumors, and from samples without visible tumors, tissue was taken from the site of tumor injection. The samples were cut into smaller pieces and incubated with Tumor Dissociation Kit (Miltenyi Biotec©) according to the manufacturer’s instructions and processed using a gentleMACS™ dissociator (Miltenyi Biotec©). After dissociation, the samples were passed through a 70 μm strainer, washed with HBSS, and myelin was removed by resuspending the samples in 25% bovine serum albumin (BSA) (Sigma-Aldrich, #A9647) and centrifuged at low brake (brake=2) for 20 minutes at 4°C. The samples were then resuspended in Hanks’ solution and passed through a 40 μm cell strainer.

### Cytometry Cell Staining

For extracellular staining, the samples were incubated with Live/dead dye (BD) for 15 minutes on ice. After washing, the cells were resuspended with CD16/CD32 (Mouse BD Fc Block™) and incubated in FACS buffer at 4°C for 15 minutes. Subsequently, the antibody mix (Table 1) was added and the samples were incubated in FACS for another 15 minutes at 4°C. The cells were then washed with FACS buffer and fixed with 2% paraformaldehyde (PFA). For intracellular staining, the cells were incubated in True-Nuclear Transcription Factor Permeabilization Buffer (#424401 Biolegend) for permeabilization. The antibody mix was prepared in perm wash and incubated for 20 minutes at room temperature. Finally, the samples were washed and resuspended in FACS buffer for analysis. Data analysis was performed using a Cytoflex LX Beckman Coulter, and the results were analyzed in Cytobank©. Manual gating was applied to examine cell populations, recording 10,000 to 20,000 events.

### Single-cell Library Preparation and Sequencing

Single-cell libraries were prepared using the BD Rhapsody™ platform (BD Biosciences) and followed as described before^59^. Naïve and TdLN single-cell suspensions were labeled sequentially using the mouse Single Cell Sample Multiplexing Kit (BD Biosciences, #633793) and BD AbSeq Ab-Oligos targeting αCD11b (#940008 M1/70), αCD200 (#940208 OX90), αF4/80 (#940131 T45-2342), and αCD103 (#940136 M290). The pooled samples were loaded onto a BD Rhapsody cartridge (BD Biosciences, #633733), and mRNA from single cells was captured using Cell Capture Beads according to the manufacturer’s protocol (BD Biosciences, #633731). Targeted cDNA libraries of genes in the BD Rhapsody Immune Response Panel Mm (BD Biosciences, #633753), supplemented with additional genes (**Table 2**), along with Sample-tag and BD AbSeqs, were prepared using the BD Rhapsody Targeted mRNA and AbSeq Amplification kit and protocol (BD Biosciences, # 633774). The final libraries quality was assessed using an Agilent 2200 TapeStation with HS D5000 ScreenTape (Agilent Technologies, Santa Clara, CA, #5067-5592), and concentrations were measured using a Qubit Fluorometer with the Qubit dsDNA HS kit (Thermo Fisher Scientific, #Q32854). The libraries were diluted to 2nM and pooled for paired-end sequencing (Read1: 51bp, Read2: 71bp + i7 Indexes: 8bp) on a NovaSeq 6000 S Prime sequencer (Illumina, San Diego, CA) at the SNP&SEQ Technology Platform (Uppsala, Sweden).

### Analysis of Single-cell RNA Sequencing Data

Fastq files from the SNP&SEQ Technology Platform were processed using the Rhapsody analysis pipeline (BD Biosciences) on Seven Bridges. Recursive Substitution Error Correction (RSEC) and Distribution-Based Error Correction (DBEC) algorithms developed by the manufacturer (BD Biosciences) were used to correct for PCR and sequencing errors. Final expression matrices contained RSEC- and DBEC-adjusted counts. Pooled samples were deconvoluted using Sample-tag reads. Multiplets were assigned if the count for two or more Sample-tag antibodies exceeded the minimum threshold. Cells that did not meet the criteria for singlets or multiplets were classified as undetermined. The final read depths for the library were as follows: mRNA 72,938.31 average reads/cell and 99.72% sequencing saturation, AbSeq 1,968.14 average reads/cell and 88.12% sequencing saturation, within the expected range of the manufacturer’s recommendations for targeted transcriptomics sequencing (BD Biosciences).

### Single-cell Sequencing Analysis Workflow

The analysis of single-cell sequencing data was performed using Partek Flow build version 10.0.21.0411 (Partek Inc., St. Louis, MI). RSEC-adjusted molecule counts were used, and undetermined cells, multiplets, and cells with <50 detected transcripts were filtered out. Data were normalized as counts per 10,000, with an offset of 1 count, and Log2 transformation was performed as recommended by the manufacturer (BD Biosciences) using the data normalization module in Partek Flow. Dimensionality reduction using PCA followed by UMAP (default settings) was performed on all genes and proteins. Clustering was performed using the Louvain clustering algorithm (default settings, nearest neighbor type: NN-Descent, number of nearest neighbors: 30), and biomarkers for each cluster were computed using ANOVA tests (default settings) with Benjamini-Hochberg correction for false discovery rate (FDR) values ≤ 0.05 and log2 fold change ≥1.5.

Sequencing was performed by the SNP&SEQ Technology Platform in Uppsala. The facility is part of the National Genomics Infrastructure (NGI) Sweden and Science for Life Laboratory. The SNP&SEQ Platform is also supported by the Swedish Research Council and the Knut and Alice Wallenberg Foundation.

## Authors contributions

OB and MH contributed to the original draft, methodology, analysis, and data curation. LH conducted the OX40 therapy *in vivo*, JN participated in the *in vivo* experiments, writing and visualization, SK performed the analysis of dendritic cells in cytometry, and MR participated in the cytometry and single-cell RNA sequencing experiments. MH provided supervision and acquired funding. All authors reviewed and approved the manuscript.

## Acknowledgements

We acknowledge Ana Dimberg and Magnus Essand form IGP Uppsala University for support in the project and data discussion. We thank our thesis worker Jenifer Lundqvist for their technical support and the BioVis Core Facility (Uppsala University) for assistance with flow cytometry. This work was supported by grants from the Cancerfonden

---

**Supple Figure 1.- Maintenance of T-Cell Profile in TdLN after *Ex-Vivo* Restimulation.** A schematic representation of the gating strategy for T-cell profiling **(A).** *Ex-vivo* experiments gating strategic **(B).** Quantification of exhausted T cells (percentage of PD-1+/TIM3+) from proliferative cells **(C)**. Graphs display mean ± SEM; analyzed using two-way ANOVA with Tukey’s correction.

**Supple figure 2.- DCs express activation markers and show reduction in the antigen-specific T cell memory response in cervical TdLN.** gating strategy for dendritic cells and activation markers (A). TdLN DCs were isolated at day 12 post-tumor inoculation. Quantification of activation markers CD86 or CD40 on DCs from SB28-OVA (left panel) or EG7-OVA tumors (right panel) (B). Percentage of DCs from the spleen at day 12 post-tumor inoculation, as well as SIINFEKL+ DCs (C). Quantification of Tetramer OVA+ regulatory T cells (CD4+ CD25+ Foxp3+) from the spleen (D). Quantification Tetramer OVA+ Effector Memory (EFM) (CD44+ CD62L-) and Central Memory (CM) (CD44+ CD62L+) form SB28-OVA (left panel) or EG7-OVA (right panel) (E). Graphs display mean ± SEM; analyzed using, two-way ANOVA with Tukey’s correction or unpaired t test.

**Supple Fig3. Cervical TdLN DCs exhibit predominant immunosuppressive activity dependent on *Batf3***. Representative dot plot showing enriched DCs **(A).** *Batf3* knockout mice were inoculated i.c. or s.c. On day 12 post-tumor inoculation, TdLN were harvested, and DCs were enriched and co-cultured with pre-stained OT cells in the presence of OVA peptide. The percentage of Tregs (CD4+ CD25+ Foxp3+) and Cytotoxic T cells CD8+ (CD107+) **(B)** was calculated. DCs from TdLN at day 12 post-inoculation were enriched. A mix of DC-c and DC-p was incubated at different concentrations and co-cultured with pre-stained OT cells in the presence of OVA peptide (C). Percentage of Tregs and cytotoxic T cells CD8+ from SB28-OVA **(D)** or EG7-OVA **(E),** as well as the percentage of progenitor-exhausting T cells from SB28-OVA **(F)** or EG7-OVA **(G)** was calculated. Unspecific peptide was used as a control. (n = 5–7 mice/group). Statistics: two-way ANOVA with Tukey’s correction.

**Supple Figure 4: OX40 promote DC response and enhanced tumor response with corticosteroid combination confers survival benefits.** Schematic Representation of Experimental Procedures (A): Mice received inoculation with SB28-OVA, followed by i.p. administration of either αOX40 or IgG Ctrl on days 4, 7, and 10 post-tumor inoculations. TdLN-c was harvested at day 12 post-tumor inoculation. The proportion of memory long-lived (CD127+ KLRG1-) and short-lived (CD127+ KLRG1+) CD4+ and CD8+ T cells (B), along with exhausted PD-1+ and TIM-3+ (C), and progenitor-exhausted (D) T cells CD4+ and CD8+, were calculated. On day 22 (E) or at the endpoint (F), the mice were euthanized, and tumor-infiltrating lymphocytes (TILs) were extracted. Proliferative (Ki67+), cytotoxic (CD107a+), exhaustive (PD-1+ and TIM-3+), progenitor exhaustion (TCF-1+ and PD-1+), and Treg profile (CD25+ FOXp3+) from mice with or without macroscopic tumors were calculated by flow cytometry. Schematic Representation of Survival Experiment (G): Mice were inoculated i.c. with SB28 parental and treated with αOX40 or IgG Ctrl as mentioned before. On day 22 post-tumor inoculation, mice received dexamethasone 2mg/Kg i.p. daily for 5 days. Kaplan-Meier survival curves of tumor-bearing mice (H). Statistical analysis was performed using the two-way ANOVA with Tukey’s correction. Key to statistics: *p < 0.05, **p < 0.01, ***p < 0.001. Bar graphs show mean ± SEM.

**Supple Figure 5. Differential scRNA-seq expression from DCs**. Heat map was generated with values presented as averaged Z scores of genes related with GESA of DCs-p versus DCs-c with significant pathway (**A, B**). (Benjamin–Hochberg adjustment p values <0.05) are shown. Dot plot representative gating of mregDC CD200, CD40 (**C**) and CD200 CCR7 (**D**). Representative Dot plot MHCII+ CD3+ of blood sample from mice Control or treated with FTY720 (**E**).

**Supplementary Table 2.**
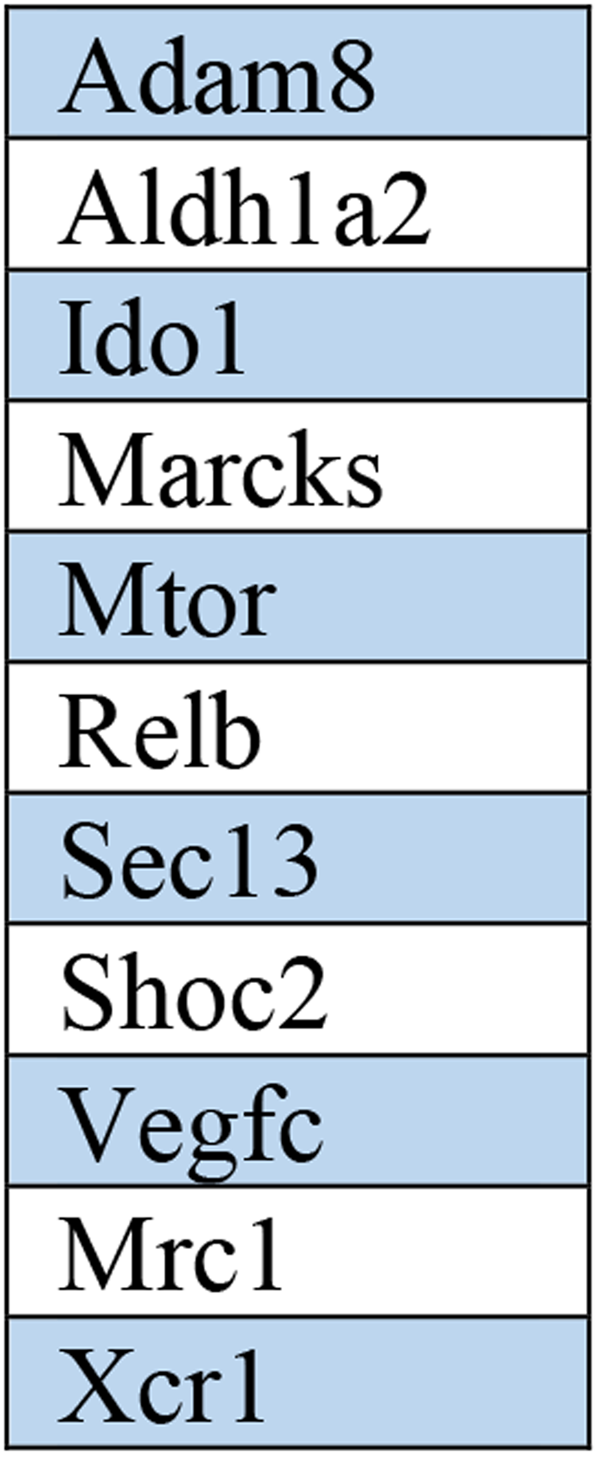
Additional genes used in the targeted transcriptomics assay, related to Figure 5 and supplementary Fig 5.

|  |
| --- |
| Adam8 |
| Aldh1a2 |
| Ido1 |
| Marcks |
| Mtor |
| Relb |
| Sec13 |
| Shoc2 |
| Vegfc |
| Mrc1 |
| Xcr1 |

