## Supplementary figures and images for "Brain tumor induce immunoregulatory dendritic cells in tumor draining lymph nodes that can be targeted by OX40"

### Supplementary figure 1

#### Supplementary Figure 1

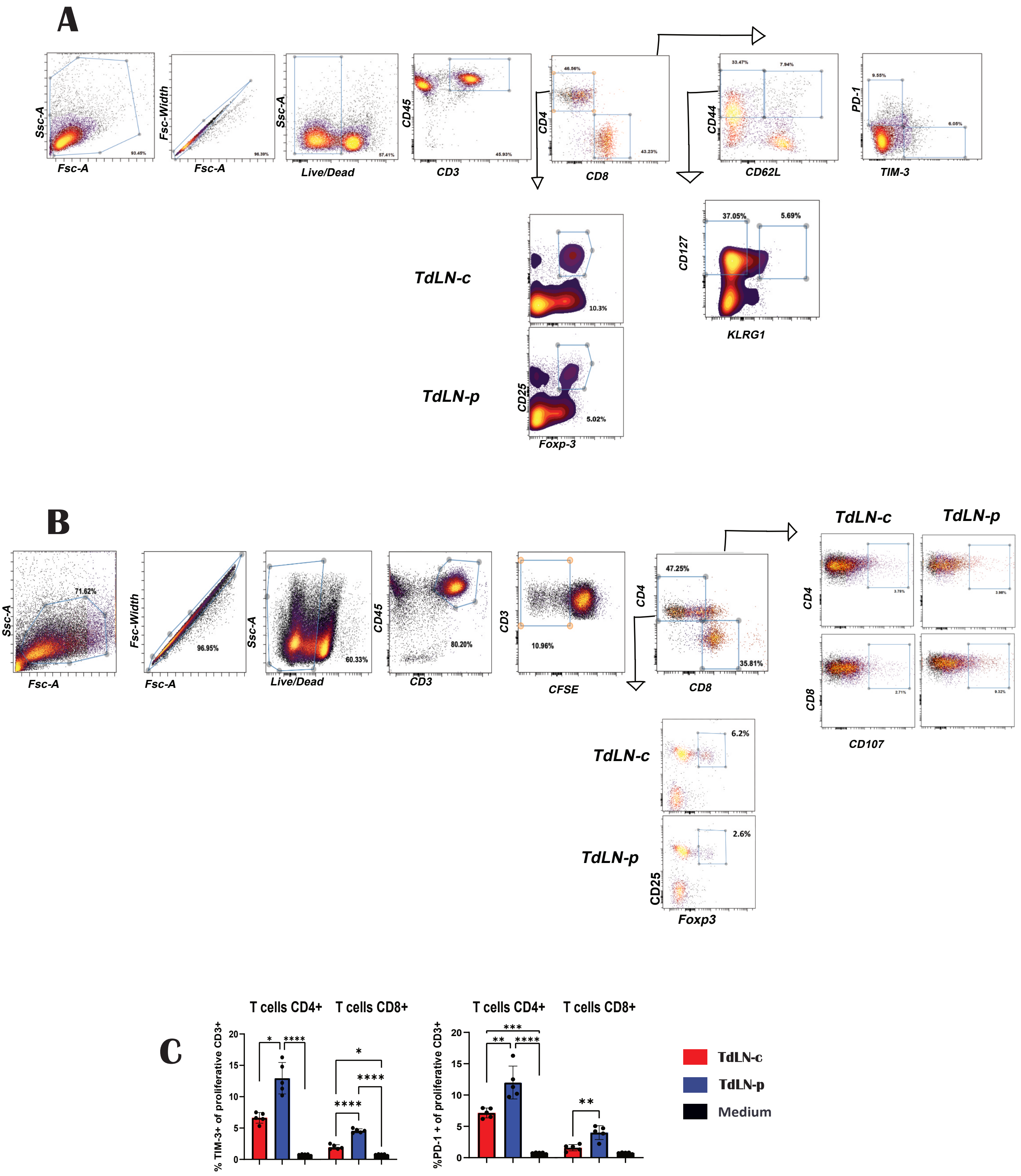

### Supplementary figure 2

#### Supplementary Figure 2

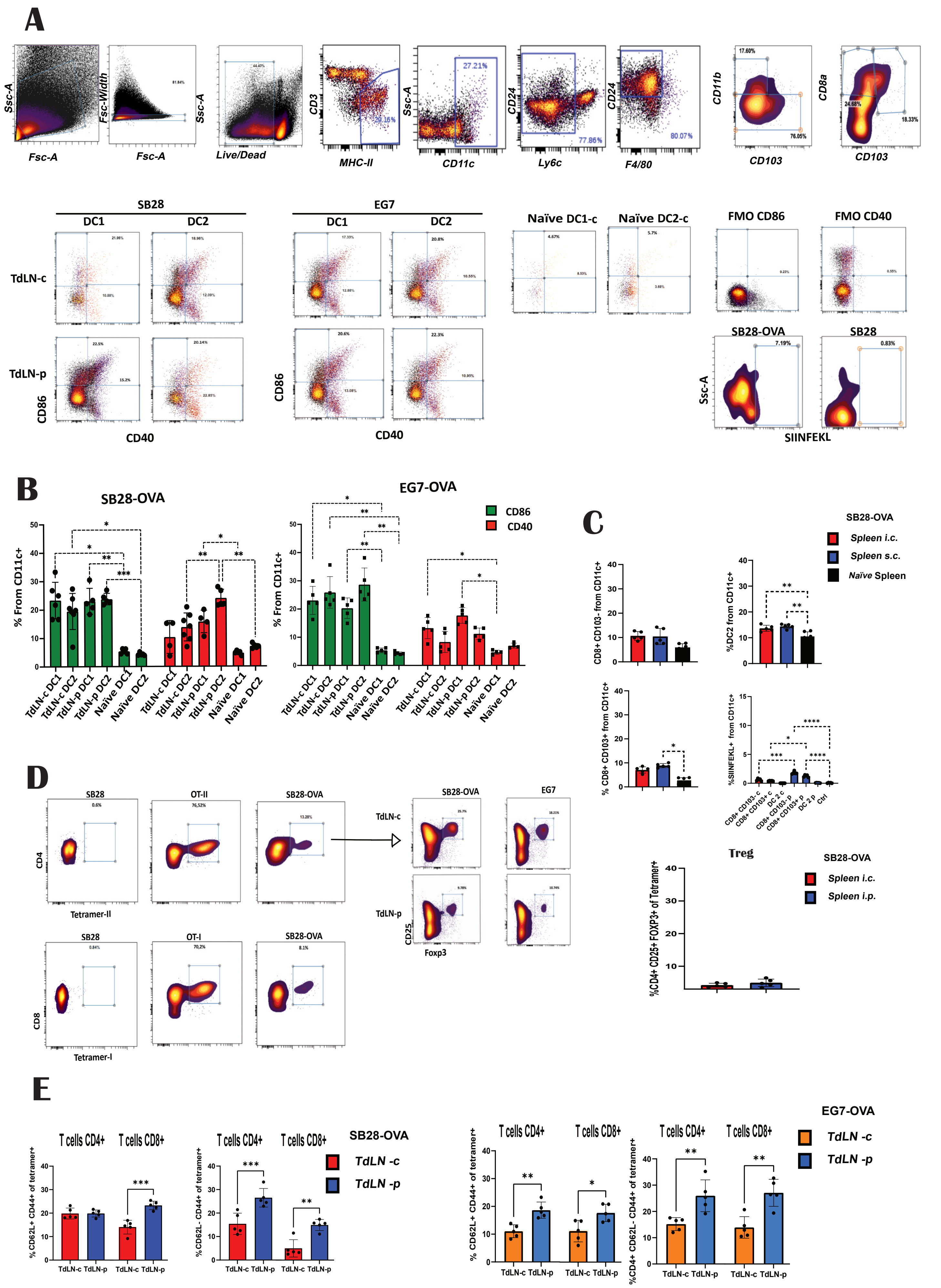

### Supplementary figure 3

#### Supplementary Figure 3

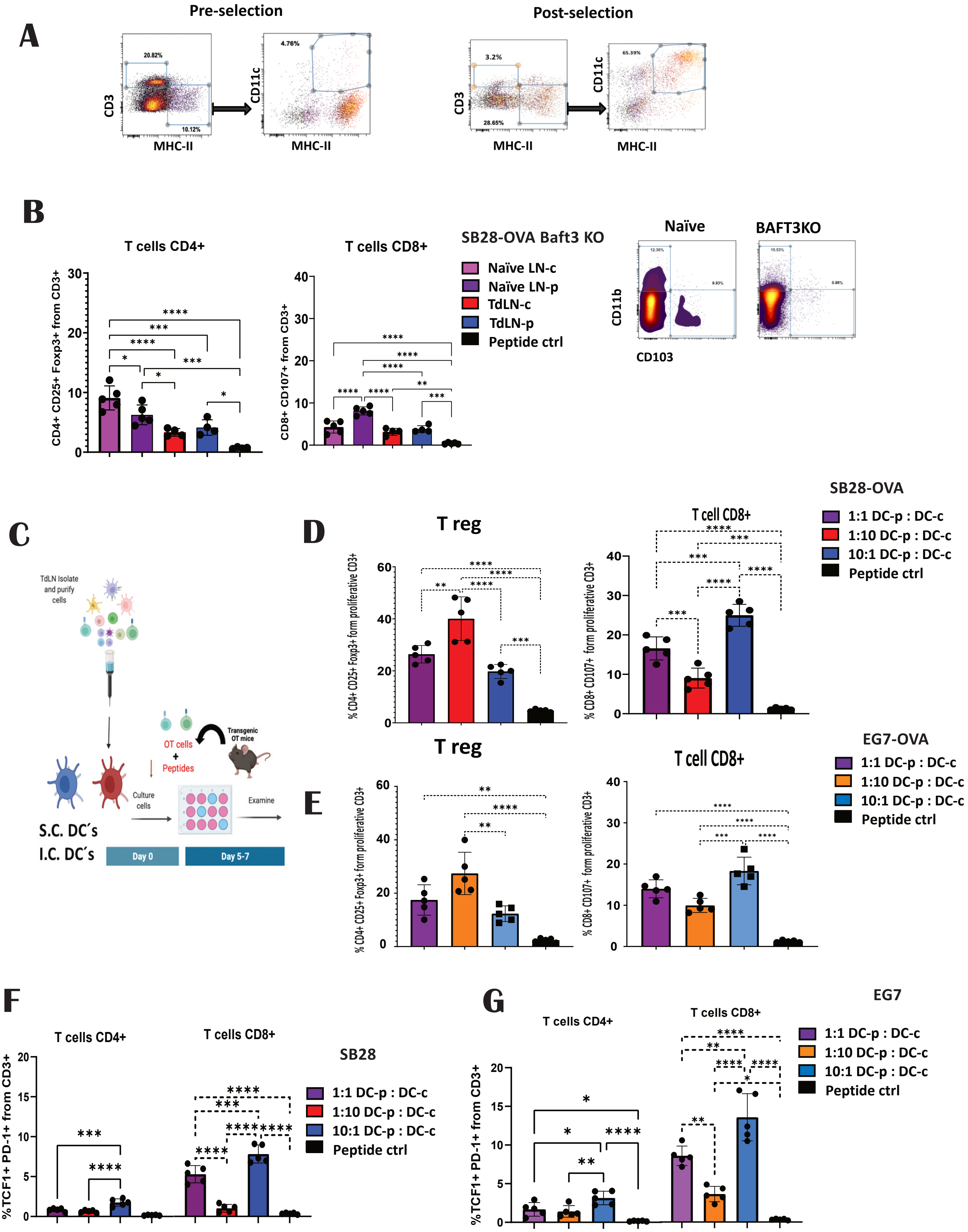

### Supplementary figure 4

#### Supplementary Figure 4

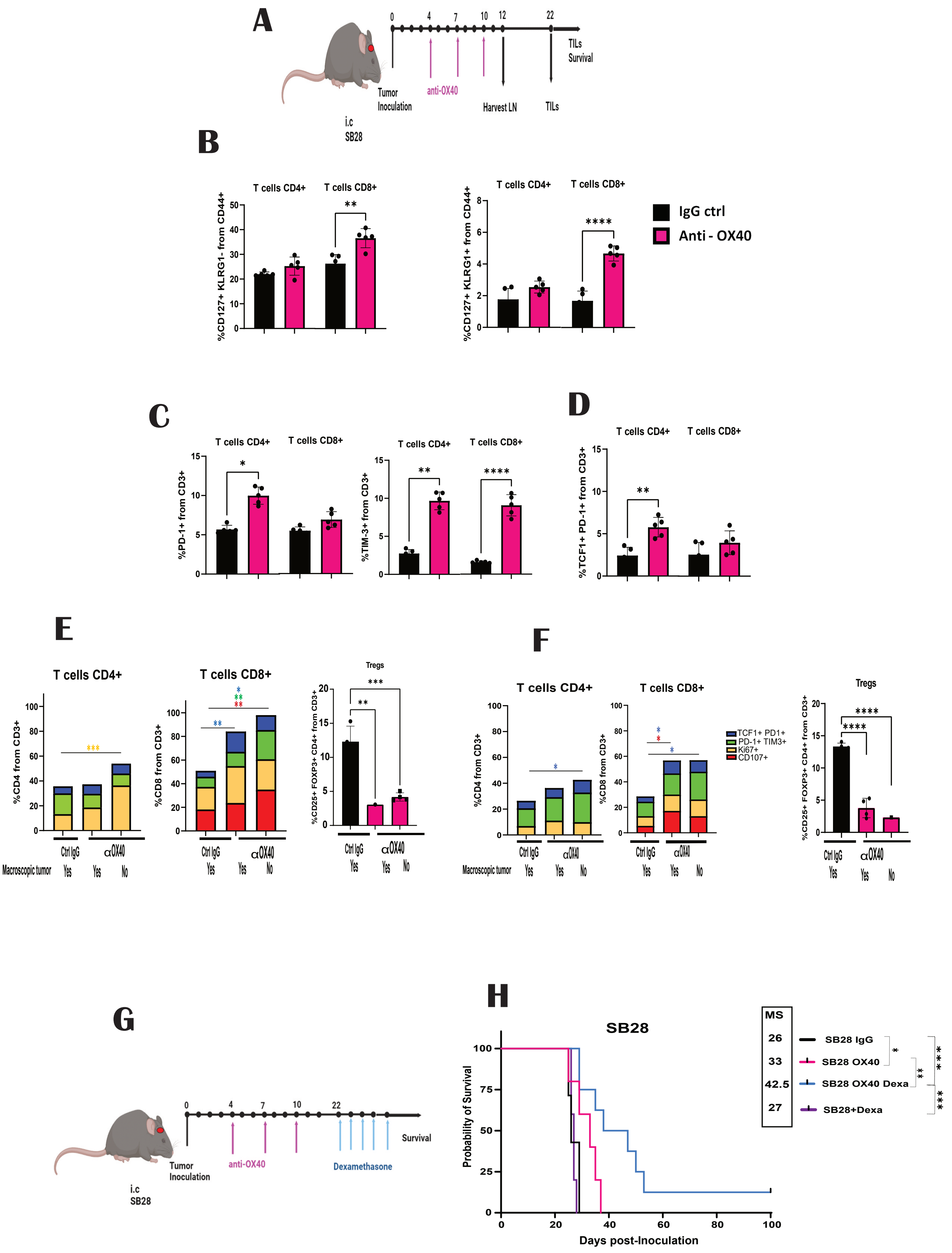

### Supplementary figure 5

#### Supplementary Figure 5

A

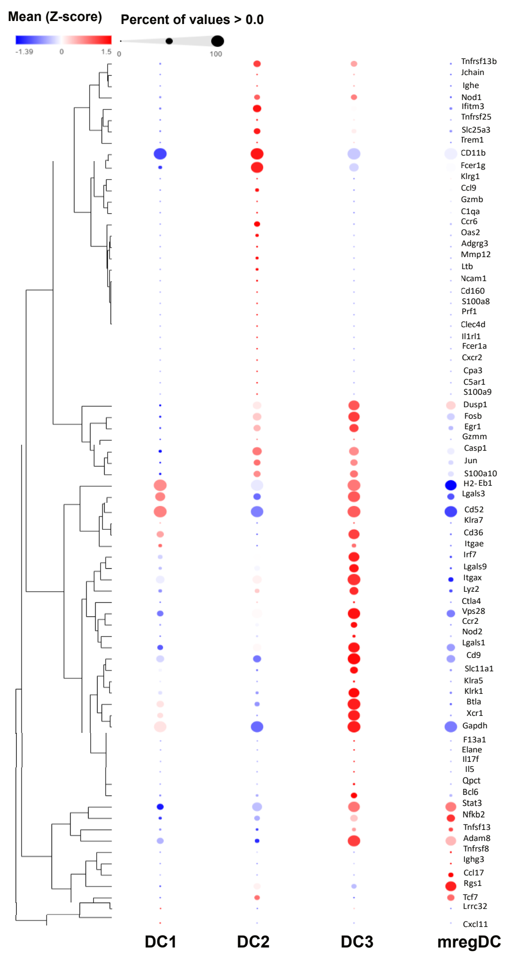


B

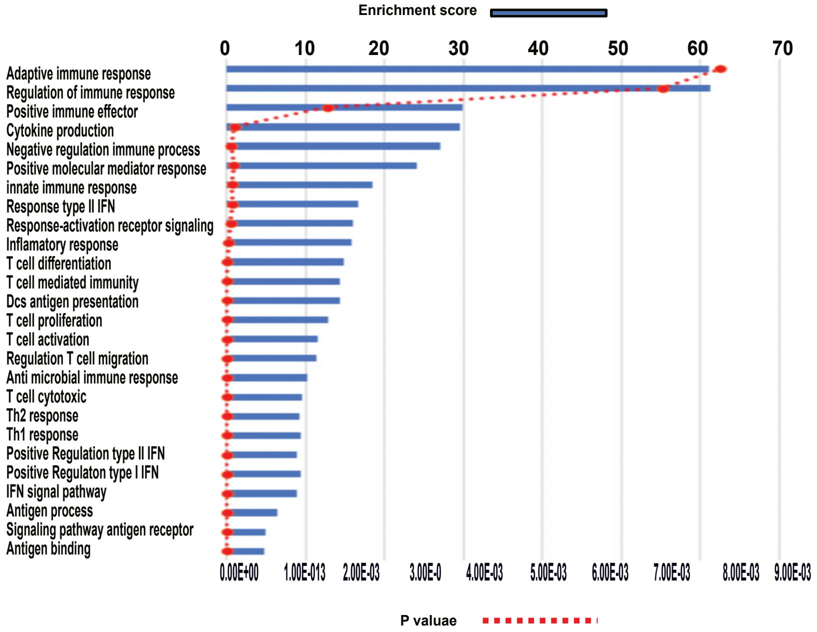


C

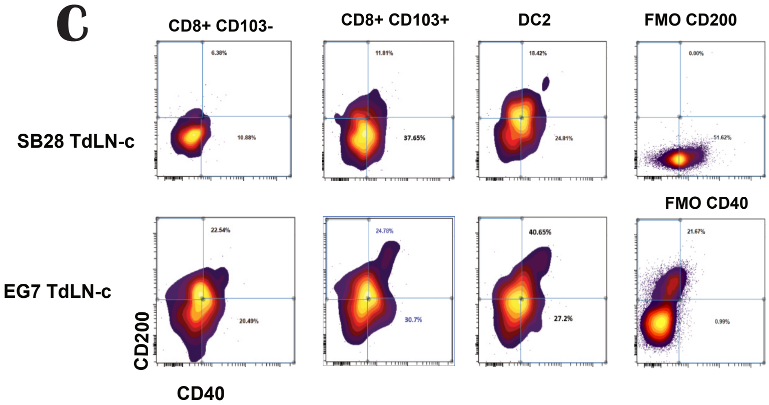


D

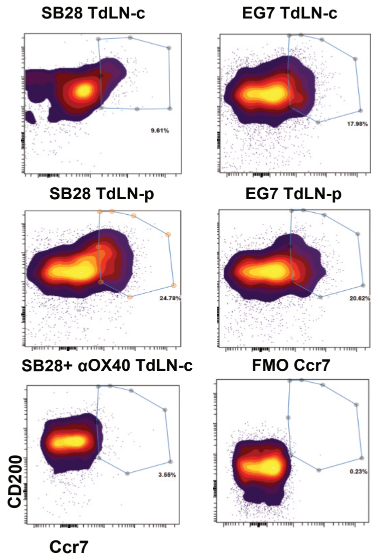


E

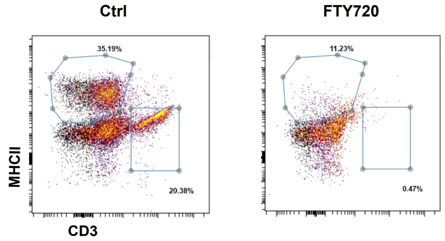
